# Neurons in the medial prefrontal cortex are involved in spatial tuning and signaling upcoming choice independently from hippocampal sharp-wave ripples

**DOI:** 10.1101/2024.09.09.610935

**Authors:** Hanna den Bakker, Fabian Kloosterman

## Abstract

The hippocampus is known to encode spatial information and reactivate experienced trajectories during sharp-wave ripple events. These events are thought to be key time-points at which information about learned trajectories is transferred to the neocortex for long-term storage. It is unclear, however, how this information may be transferred and integrated in downstream cortical regions. In this study, we performed high-density probe recordings across the full depth of the medial prefrontal cortex and in the hippocampus simultaneously in rats while they were performing a task of spatial navigation. We find that neurons in the medial prefrontal cortex encode spatial information and reliably predict upcoming choice on a maze, and we find that a subset of neurons in the mPFC is modulated by hippocampal sharp-wave ripples. However, sharp-wave ripple modulation does not appear to be the main driving factor in predicting upcoming choice. This indicates that the integration of spatial information requires the collaboration of different specialized populations of neurons.

## INTRODUCTION

The hippocampus is known to be important for spatial navigation. This is largely attributed to the spatially specific firing patterns of place cells, which fire sequentially as animals navigate through an environment in the timeframe of theta frequency (∼8Hz) oscillations. These sequential firing patterns are often repeated or *‘‘*replayed*’’* during short high frequency oscillations called sharp-wave ripples (SWRs) while the animal is in a state of behavioral immobility or sleep. The function of these replay patterns is still somewhat under debate, but during awake immobility periods they have been proposed to either predict immediate future behavior^1–5^, or reflect specific past experience^6^, and adapt to reward^7,8^ and task demand^9^, while during sleep, they are thought to be important for memory consolidation^10^.

A running hypothesis is that the hippocampus transfers information to neocortical areas during these SWR events and this is thought to be a crucial mechanism to facilitate spatial learning^11^. Congruently, it has been shown that a subset of neurons in the medial prefrontal cortex shows modulated firing patterns following SWRs in the hippocampus, where some neurons show an increase and others a decrease of firing rate, both in a 200 millisecond window following SWR onset^12,13^. Disrupting this window of activity in the mPFC slowed the acquisition of a spatial rule reversal^14^.

The mPFC can be divided into three regions: the anterior cingulate cortex (ACC), the prelimbic cortex (PLC) and the infralimbic cortex (ILC), which have been shown to have distinct functions in the context of fear-related memory^15–17^, cognitive flexibility^18^ and preparatory attention and error-related events^19–23^. However, due to practical limitations related to the anatomical layout of the region, it has been challenging to study electrophysiological differences between all three subregions simultaneously with sufficient single cell data to draw meaningful conclusions about functional specializations. Some authors argue that the mPFC subregions are not distinguishable as three separate subunits, but rather function as a continuum^24^. Especially the function of the PLC is under debate, and a recent study suggested to divide the PLC further into a dorsal and ventral section, where the dorsal PLC was found to be more related to ACC firing patterns, and the ventral PLC more closely resembled the ILC^25^.

The response to SWRs in the medial prefrontal cortex has been found to be specific to the place cell activity in the hippocampus during the SWR, where non-local replay events in the hippocampus are followed by a higher firing rate modulation in the medial prefrontal cortex, and this preference is predictive of future behavior^13^. The modulation to SWRs also seems to be stronger when the SWRs occur during behavioral immobility or rest, compared to sleep SWRs^26^, potentially indicating a different mechanism underlying the two states.

Cells in the mPFC that are modulated by SWRs, are more likely to be phase-locked to the hippocampal theta rhythm seen mostly while the animal is actively exploring an environment^12,13^. Ripple and theta oscillations are independent brain states, occurring primarily during rest and active exploration, respectively. Therefore it is interesting that mPFC neurons are modulated by both states, which suggests a network of prefrontal-hippocampal activity that is co-modulated during both rest and active exploration^13^.

Although the spatial specificity is lower in mPFC neurons compared to the hippocampus, some papers have reported replay trajectory events to occur in the mPFC as well^27–31^. The coordination with replay events in the hippocampus remains unclear, where one study reports coordinated replay trajectory events between the hippocampus and mPFC^31^, while another study found that mPFC replay occurs independently of hippocampal replay^28^. During sleep it has been found that reactivation of previous experiences in the mPFC does occur time locked to SWRs in the hippocampus, but only when a spatial rule on the maze had been learned^30^, indicating that this interaction may be selective depending on the context in which it occurs. It has also been found that the mPFC engages in replay events particularly after error trials^29^, which may reflect a mechanism of negative feedback during learning. Finally, cells in the mPFC that are modulated by SWRs in the hippocampus have been shown to be less tuned to space^12^, and therefore perhaps this modulation of cellular activity is not necessarily (only) a reflection of spatial replay trajectories, but may reflect another non-spatial mechanism instead.

In this study, we performed high-density probe recordings across the full depth of the mPFC and in the hippocampus simultaneously while animals are performing a behavioral task of spatial navigation, to shed light on firing properties of SWR-modulated and unmodulated cells in the mPFC. We find that mPFC cells that are not modulated by hippocampal SWRs seem to be active at key time points in spatial decision making.

## Results

### Chronic Neuropixels probe recordings across mPFC subregions

Adult Long Evans rats were implanted with either two Neuropixels 1.0 probes (mPFC and hippocampus; N=5 animals) or a single probe (mPFC; N=1 animal; see Methods and Fig. 1a-b and Supplementary Table 1). Signals from the mPFC were classified as originating from the dorsal or ventral subregion based on the implantation depth of the electrodes and the histological verification of the probe track (see Methods and Supplementary Fig. S1). Both subregions showed ample spiking activity (for example see Fig. 1b). In five animals, data from both dorsal and ventral mPFC was obtained and in one animal only data for the dorsal mPFC was included (due to the probe implantation being too posterior; see Supplementary Fig. S1). Spikes were assigned to a cluster (putative neuron) using automated spike sorting followed by manual curation (see Methods), and up to 203 neurons were isolated from a single subregion per session (see Fig. 1d and Supplementary Table 1). For all analyses presented here, data was averaged per session, so that sessions with a high number of neurons did not skew the results. Due to the light-weight and compact design of the implant^32^, animals were able to move freely on an elevated track. While we recorded neural activity, the animals performed a spatial rule switching task in which they learned 4 different rules over the course of multiple days (see Methods and Fig. 1c). Rules were based on either allocentric (*“*go North*”* or *“*go South*”*) or cue-based (*“*go to illuminated arm*”* or *“*go to dark arm*”*) strategies, which were to be deducted by the animals through trial and error. Each session started with a period of 20 minutes rest in a dark resting box. After this, the animal was placed by the experimenter pseudo-randomly on either the East or the West arm of the maze and it had to learn to go (1) to the North arm, (2) to the South arm, (3) to the illuminated arm (either South or North) or (4) to the non-illuminated (South or North) arm. The order of the rules was counterbalanced between animals. The illumination of the arm was always present (also during allocentric learning) and was pseudo-randomly determined for each trial. After the animal made its choice, it was picked up by the experimenter, and placed back in the dark resting box for 30-60 seconds before continuing with the next trial. The animals were trained for a maximum of 1.5 hours or 60 trials per day. At the end of the session, animals were placed back in the dark resting box for up to 60 minutes.

**Figure 1.**
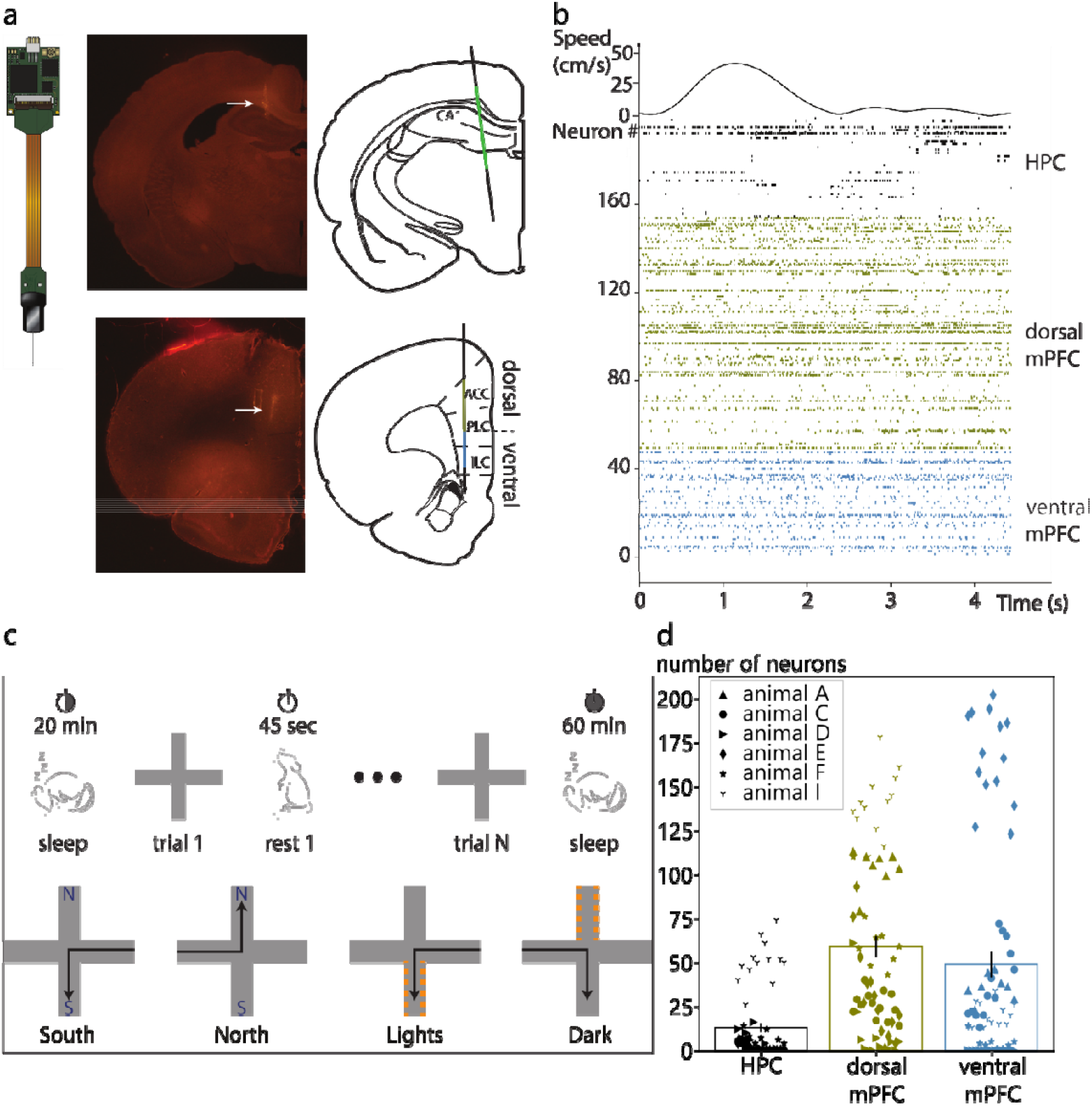
Chronic Neuropixels probe recordings across mPFC subregions. **(a)** Left: an image of the Neuropixels probe, right: the probe mapping from one example animal where a part of the trace is seen going through the hippocampus (top) and mPFC (bottom). **(b)** An example segment of data with the animals running speed (top) and spiking activity recorded from the hippocampus, dorsal mPFC and ventral mPFC (top to bottom). **(c)** An overview of the behavioral task. Each day starts with a period of rest, followed by rule learning in the maze. Each trial is interleaved with a short period (∼45 seconds) of rest, and the day ends with a period of rest or sleep of up to 60 minutes. Four different rules have to be learned over the course of multiple days, based on allocentric and cue-based strategies. **(d)** The number of sorted neurons per session for each (sub-)region. The different markers indicate different animals.

### Neurons in the dorsal and ventral mPFC are differentially modulated by hippocampal SWRs and theta phase

The mPFC is known to contain functionally distinct cell populations that differ in the modulation of their activity by hippocampal SWRs and theta oscillations^12^. Both SWR-excited and SWR-inhibited cell populations (but not SWR unmodulated cells) have been shown in the past to be phase-locked to hippocampal theta oscillations and they are more likely to co-fire with cell members of their own group^12^. Here we test if these cell populations are equally present across the dorsal-ventral extent of the mPFC.

We first confirmed the presence of SWR and theta-modulated subpopulations of mPFC neurons while animals were on the maze. Following hippocampal SWR onset, 13.96% of all mPFC neurons significantly increased their firing rate (SWR-excited) and 5.46% of neurons decreased their firing rate (SWR-inhibited). The majority of neurons were not modulated by hippocampal SWRs (80.59%; SWR-unmodulated, see Fig. 2a-d). These neurons had a lower overall firing rate than SWR modulated neurons (mean firing rates in Hertz, SWR-excited: M=6.25, SEM=0.49, SWR-inhibited: M=9.31, SEM=1.00, SWR-unmodulated: M=3.87, SEM=0.33, ANOVA: F(142)=19.57, p<.001, see Fig. 2e). Cells that changed their firing rate following hippocampal SWRs also were more likely to be phase-locked to the hippocampal theta oscillation. Overall, 38.46% of cells in the mPFC and 79.42% of cells in the hippocampus showed significant theta phase locking, similarly to previously reported percentages^33^. Theta phase-locking was more prevalent among the population of SWR-modulated mPFC cells (M=55.40%, SEM=4.50) than the population of SWR-unmodulated cells (M=33.53%, SEM=1.94, Mann-Whitney U test: U(104)=1993, p<.001; see Fig. 2f-g). These data are consistent with the idea that SWR- and theta-modulated cells form a functional subpopulation in the mPFC.

**Figure 2.**
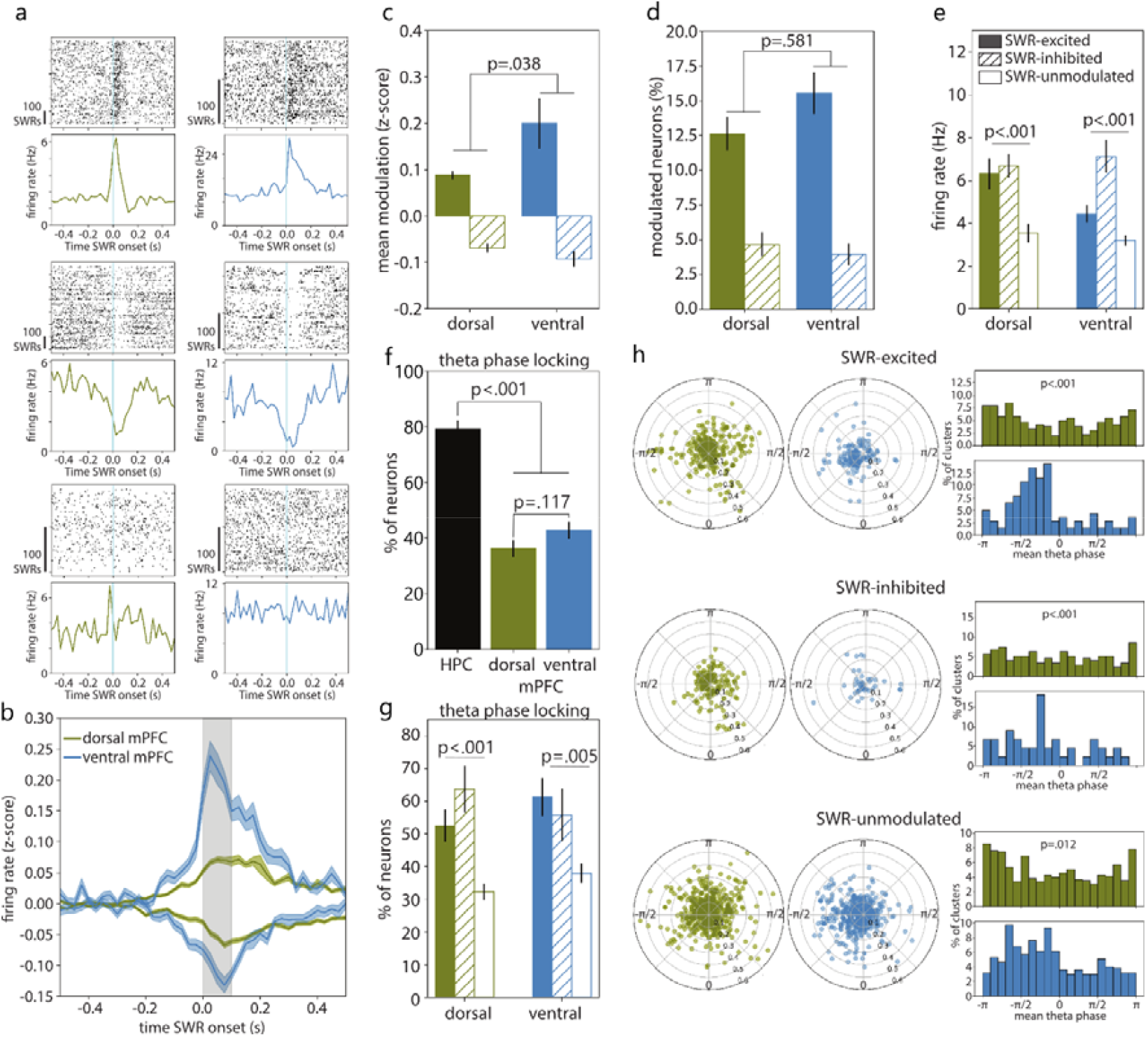
Neurons in the dorsal and ventral mPFC are differentially modulated by hippocampal SWRs and theta phase. **(a)** Six example neurons in the mPFC that are SWR-excited (top), SWR-inhibited (middle) and SWR-unmodulated (bottom; dorsal mPFC in olive, ventral mPFC in blue). **(b)** The average normalized firing rates around SWR onset of all SWR-excited and SWR-inhibited neurons in the dorsal and ventral mPFC. Data is presented as mean of all neurons ± SEM. **(c)** The mean of the SWR modulation per subregion during periods of exploration in the maze. The p-value indicates the result of an independent samples t-test between the absolute modulation scores in the dorsal versus ventral mPFC. **(d)** The percentage of neurons that is SWR-excited (solid fill) and SWR-inhibited (dashed fill) per mPFC subregion. The p-value indicates the result of an independent samples t-test between the dorsal and ventral mPFC. **(e)** The mean overall firing rates per subregion of SWR-excited (solid fill), SWR-inhibited (dashed fill) and SWR-unmodulated (no fill) neurons in the maze. The p-values indicate an ANOVA between the different SWR modulation categories. **(f)** The percentage of neurons that is phase-locked to hippocampal theta per brain region. The p-values indicate the results of Mann-Whitney U test between the hippocampus and mPFC, and between the dorsal and ventral mPFC. **(g)** The percentage of neurons that is phase-locked to hippocampal theta, divided between mPFC neurons that are SWR-excited (solid fill), SWR-inhibited (dashed fill) and SWR-unmodulated (no fill). The p-values indicate an ANOVA between the different SWR modulation categories. **(h)** Left panels: the mean angle (-π to π) and (0.0 to 0.6) for all dorsal mPFC clusters that were significantly phase locked to hippocampal theta, divided between SWR-excited clusters (top), SWR-inhibited clusters (middle) and SWR-unmodulated clusters (bottom). Middle panels: the mean angle and for all phase locked clusters in the same configuration as the left panels, but for the ventral mPFC. Right panels: histograms of the mean angles of all theta phase-locked clusters that were SWR-excited, SWR-inhibited or SWR-unmodulated (top to bottom). The p-values indicate the result of a Kolmogorov-Smirnov test to test the difference in distribution between the dorsal (olive) and ventral (blue) mPFC.

Next, we tested if and how the SWR- and theta-modulated cell populations vary in the dorsal and ventral subregions of the mPFC. We observed similar percentages of SWR-excited and SWR-inhibited neurons in both subregions. However, the activity of SWR-excited and SWR-inhibited neurons in ventral mPFC was more strongly modulated by SWRs than the activity of neurons in the dorsal mPFC (mean modulation absolute z-score, dorsal: M=0.09, SEM=0.01, ventral: M=0.20, SEM=0.05; independent samples t-test: t(86)=-2.11, p=.038; see Fig. 2b and Fig. 2c). The theta-modulated cells were also similarly distributed in the dorsal and ventral mPFC (percent of theta modulated neurons, dorsal: M=36.67%, SEM=3.07, ventral: M=42.88%, SEM=3.18; Mann-Whitney U test: U(104)=1139, p=.117, see Fig. 2f). In both brain regions, there was a higher proportion of theta modulated cells among the SWR-modulated compared to the SWR-unmodulated cells (percent of theta modulated neurons, dorsal, SWR-modulated: M=55.53, SEM=4.85, SWR-unmodulated: M=31.93, SEM=2.51; Mann-Whitney U test: U(98)=1794, p<.001. Ventral, SWR-modulated: M=59.14, SEM=5.63, SWR-unmodulated: M=37.46, SEM=2.96; Mann-Whitney U test: U(88)=1354, p=.005, see Fig. 2g). The preference for the hippocampal theta phase, however, was different between the dorsal and ventral subregions. Phase-locked neurons in the dorsal mPFC had a bias for firing just before the peak of the oscillation, while phase-locked neurons in the ventral mPFC were more likely to fire around the trough of the oscillation (theta phase locking dorsal mPFC versus ventral mPFC, Kolmogorov-Smirnov difference in distributions, SWR-modulated neurons: K=0.8, p<.001; SWR-unmodulated neurons: K=0.5, p=.012; see Fig. 2h).

### SWR-unmodulated neurons are spatially tuned, directionally selective and show theta cycle skipping

Previous research has demonstrated that neurons in the mPFC encode spatial information^27–29,34^, and suggested that SWR-unmodulated neurons show lower spatial coverage (i.e. higher spatial tuning) than SWR-excited and SWR-inhibited neurons^12^. Another paper found that the dorsal mPFC has higher spatial tuning properties than the ventral mPFC^35^, from which the authors concluded that spatial tuning in the mPFC is only partly dependent on direct input from the hippocampus. Here we tested whether the difference in spatial tuning between SWR-modulated and SWR-unmodulated neurons holds up when comparing the dorsal and ventral mPFC. We calculated the spatial specificity score for each cluster (see Methods) and found a higher spatial tuning score in the hippocampus (spatial information in bits/spike, M=0.64, SEM=0.06) than in the mPFC neurons (M=0.31, SEM=0.01, Mann-Whitney U test: U(106)=2129, p<.001; Fig. 3a-b), as expected from previous literature^12,28^. In contrast to the mentioned study^35^, we did not observe differences between spatial tuning properties of neurons in the dorsal and ventral mPFC (spatial information in bits/spike, dorsal: M=0.32, SEM=0.01, ventral: M=0.34, SEM=0.02; Mann-Whitney U test: U(121)=1969, p=.639). We did, however, find that SWR-unmodulated mPFC neurons showed a higher spatial tuning score (M=0.33, SEM=0.02) compared to SWR-excited (M=0.22, SEM=0.01) and SWR-inhibited neurons (M=0.11, SEM=0.01; Kruskal-Wallis: H(142)=70.52, p<.001; Fig. 3b). This effect is robust and was seen for both the dorsal and ventral subregions.

**Figure 3.**
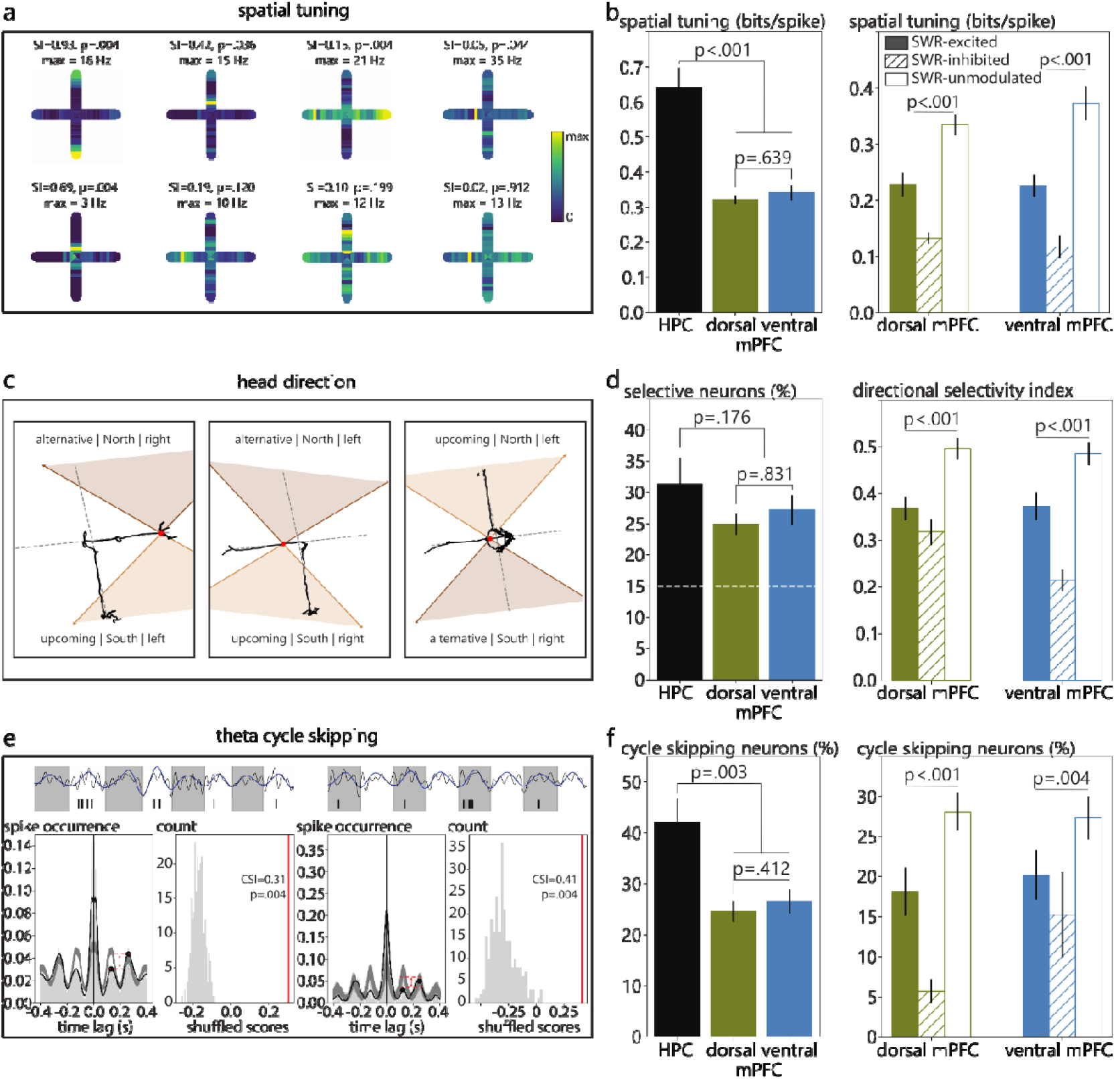
SWR-unmodulated neurons are spatially tuned, directionally selective and show theta cycle skipping. **(a)** The firing patterns of eight example neurons in the maze, showing a range of spatial tuning scores (in bits/spike) sorted from highest to lowest. **(b)** Left panel: the spatial tuning score per subregion (p-values indicate a Mann-Whitney U test). Right panel: the spatial tuning score per subregion divided between mPFC neurons that are SWR-excited (solid fill), SWR-inhibited (dashed fill) and SWR-unmodulated (no fill; p-values indicate a Kruskal-Wallis test). **(c)** Three example trials where quadrants on the maze are categorized as upcoming or alternative choice, North or South, and left or right. **(d)** Left panel: the percentage of neurons that fire significantly higher than the shuffled population for at least one of the head direction categories. The chance level is indicated at 15%, as an alpha of 0.05 was maintained for each of the three direction categories (future versus alternative, North versus South, and left versus right; p-values indicate the results of a Mann-Whitney U test). Right panel: the absolute maximum directional selectivity index per subregion for mPFC neurons that are SWR-excited (solid fill), SWR-inhibited (dashed fill) and SWR-unmodulated (no fill; p-values indicate a Kruskal-Wallis test). **(e)** Spike time auto-correlations of two example neurons where the peak of the smoothed signal (in black) at 250 ms is subtracted from the peak at 125 ms and normalized to the highest of the two peaks to obtain the cycle skipping index (left panels). To determine the statistical significance, the score is compared to a shuffled population (right panels). **(f)** Left panel: the percentage of neurons that showed significant theta cycle skipping properties per subregion (p-values indicate a Mann-Whitney U test). Right panel: the theta cycle skipping index per subregion divided between mPFC neurons that are SWR-excited (solid fill), SWR-inhibited (dashed fill) and SWR-unmodulated (no fill; p-values indicate a Kruskal-Wallis test).

Cells that fire selectively during particular head directions have been found in a variety of brain regions as well^36–38^. We next investigated whether neurons in the mPFC are linked to certain head directions and whether there is also a differential involvement of SWR-modulated and unmodulated neurons in head direction selective firing patterns. We categorized the animals’ head directions at the start of the maze in three categories (towards future choice or alternative choice, towards North or South, and towards left or right, see Methods and Fig. 3c). First, we determined for each neuron whether it spiked significantly more for one direction compared to its opposite counter part (e.g. future choice versus alternative choice). In the hippocampus 31.35% of clusters (SEM=4.20) showed significant directional selectivity, compared to 24.78% of clusters in the mPFC (SEM=1.52, Mann-Whitney U test: U(103)=1505, p=.176; chance level is 15%; see Fig. 3d). Next, for each neuron a directional selectivity index was calculated, where the difference between the firing rates within one category were subtracted from the summed firing rate of that category (e.g. the difference between firing rates during head directions towards the future choice and alternative choice, divided by the sum of both direction; see Methods). This index thus indicates the preference of each neuron to fire selectively during one type of head direction. For all three head direction categories (future versus alternative, North versus South, and left versus right), the SWR-unmodulated neurons showed significantly higher directional selectivity scores than the SWR-excited and SWR-inhibited neurons (maximum absolute directional selectivity index, SWR-excited: M=0.37, SEM=0.02, SWR-inhibited: M=0.27, SEM=0.02, SWR-unmodulated: M=0.48, SEM=0.02; Kruskal-Wallis: H(138)=41.26, p<.001, see Fig. 3f). On a population level, there was no preference for any one of the categories over another (see Supplementary Fig. S3b).

Furthermore, both the mPFC and hippocampus are known to encode information about upcoming trajectories in alternating cycles of theta oscillations^39–41^. A subset of cells in the hippocampus and mPFC are found to skip every other cycle within the theta rhythm, and neurons in the mPFC are thereby thought to favor the true upcoming trajectory over alternative choices^39^. Here we were interested to see whether we observe differences in theta cycle skipping properties in SWR-modulated and unmodulated neurons in the dorsal and ventral mPFC. For each neuron we determined whether or not it showed cycle skipping properties by looking at the probability of spiking at 125 ms (one theta cycle) and 250 ms (two theta cycles) in the spike time autocorrelation (see Methods and Fig. 3e). A subset of neurons in the hippocampus (M=42.03%, SEM=4.70) and in the mPFC (M=25.69%, SEM=1.87) showed theta cycle skipping properties. The percentage of neurons showing theta cycle skipping properties was significantly higher for SWR-unmodulated neurons, compared to SWR-excited and SWR-inhibited neurons (percentage of neurons showing theta cycle skipping properties, SWR-excited: M=22.72, SEM=2.01, SWR-inhibited: M=12.39, SEM=3.04, SWR-unmodulated: M=28.94, SEM=1.99; Kruskal-Wallis: H(140)=31.25, p<.001, see Fig. 3f and Supplementary Fig. S3c).

Together these results indicate that the population of SWR-unmodulated neurons seem to have distinct functions in terms of spatial mapping, head direction selectivity, and cycle skipping properties, with a stronger involvement compared to neurons that are either excited or inhibited by hippocampal SWRs.

### Non-local representations in the mPFC are predictive of the animals’ choice

Previous research has shown that the ensemble firing patterns in the mPFC can be used to decode spatial locations^29^. Furthermore, decoded position data from the mPFC has been found to contain excursions to non-local positions in the maze, which are thought to be related to the neurons’ encoding of choice evaluation^29^. We were interested to see whether non-local representations in the mPFC could be linked to the future arm choice of the animals as well, and whether there is a link with hippocampal SWRs and SWR-modulated firing patterns. We decoded spatial information from spiking activity along the depth of the mPFC, using a Bayesian decoding model to associate spike features with the location in the maze at which the spikes occurred (see Methods). Using these decoding models, we were able to decode the current location of the animal based on spiking activity in the mPFC accurately (median cross-validated decoding error less than 20 cm) for a total of 40 sessions from 5 different animals. We decoded the spatial information at a theta time scale (50 ms bins with 80% overlap), and observed, aside from the local representations of the current arm, segments of representations of the other arms of the maze, that were either predictive of the upcoming choice, or of the alternative choice on the other side of the maze (see methods and Fig. 4a). Segments of non-local representations were defined as any number of consecutive time bins where the summed decoded probability of one of the two goal arms was higher than 0.5 while the animal was at the start or center of the maze (see Methods). Using the difference between the sum of decoded probabilities of the segments of the upcoming choice and the segments of the alternative choice, we obtained a prediction index between 1 and -1 corresponding to the direction the animal did and did not turn, respectively. These scores could accurately predict the turning direction of the animals, while the rule was still being learned as well as when it was already well known (see Fig. 4b).

**Figure 4.**
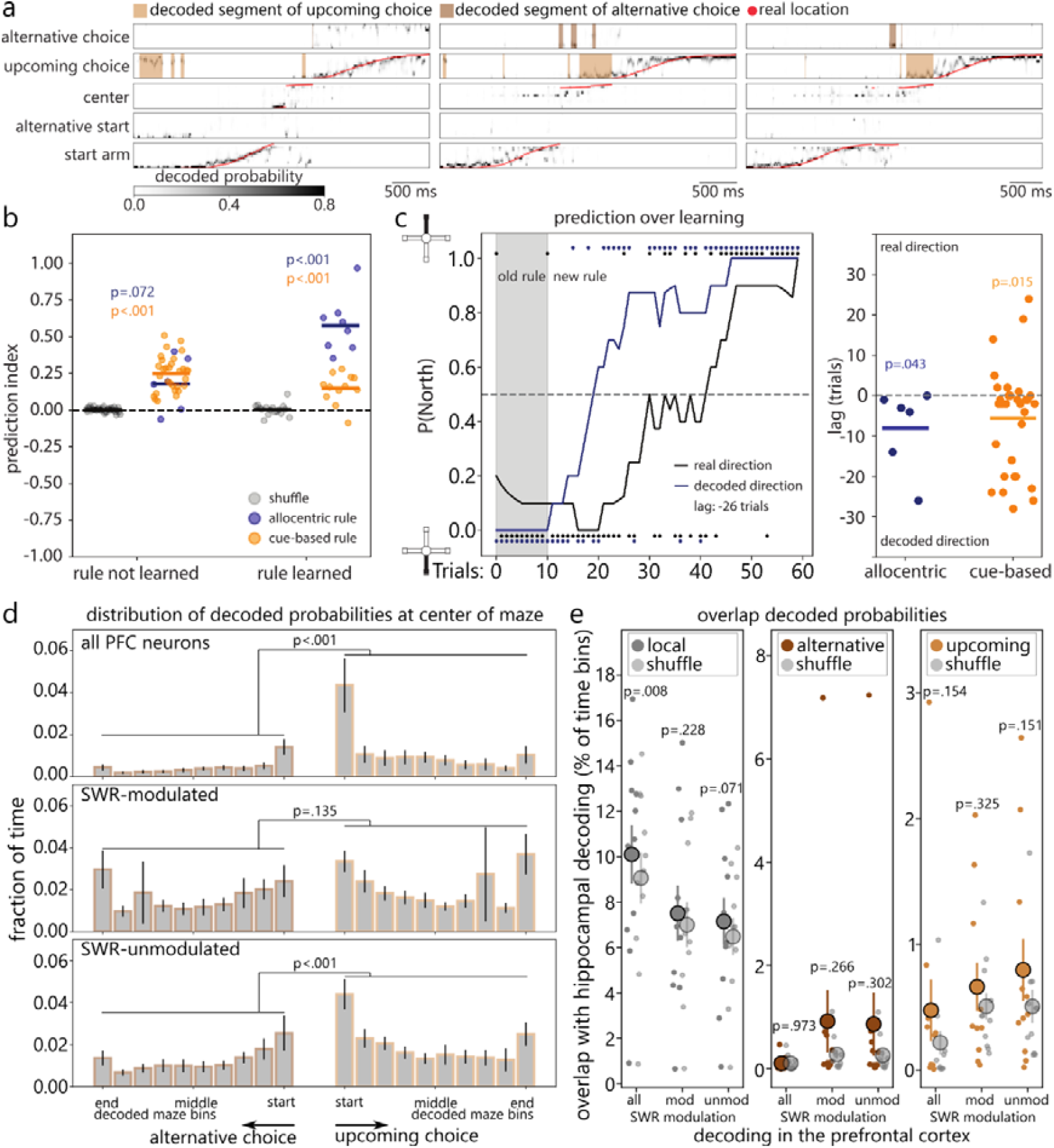
Non-local representations in the mPFC are predictive of the animals’ upcoming choice. **(a)** Three examples of trials where the decoded positions (grey scale) and real positions (red) are shown per maze segment. Segments of upcoming and alternative choice are highlighted with light and dark brown spans. **(b)** The prediction index per session based on the decoded non-local representations while the animal was at the start and center of the maze, separated for when the rule was not yet learned and when the rule was learned, and for the allocentric (navy) and cue-based (orange) rules. Shown in light grey is the prediction accuracy based on the shuffled decoded posteriors, and p-values indicate the result of a paired samples t-test between the real and shuffled predictions. **(c)** Left panel: an example learning curve (in black) with a rule switch from South to North after 10 trials, and the predicted directions based on the non-local representations overlayed (navy). Right panel: the lag (in number of trials) between the real direction and decoded direction per session for allocentric (navy) and cue-based (orange) strategies. A negative lag indicates that the decoded direction preceded the real direction over the course of learning. P-values indicate the result of a one-sample Wilcoxon test to test the deviation from zero. **(d)** The distribution of decoded probabilities at the center of the maze for all mPFC neurons (top panel), SWR-modulated neurons (middle panel) and SWR-unmodulated neurons (bottom panel). P-values indicate the results of a two-way repeated measures ANOVA between the maze bins and prediction categories (alternative versus upcoming choice). **(e)** The overlap between representations in the hippocampus and mPFC (all neurons, SWR-modulated neurons and SWR-unmodulated neurons) for the local representations (left panel), the representations of alternative choice (middle panel), and the upcoming choice (right panel). P-values indicate the results of a paired samples t-test of the percentage of overlapping decoded time bins for each session compared to the overlap of the shuffled data for that session.

Next, we asked if the predicted directions based on decoded non-local representations either preceded or followed the change in behavior. For this, we overlayed the learning curves of the animals with curves made from the predicted directions based on the non-local representations in the mPFC and performed cross-correlations between the two curves (see Methods and Fig. 4c). It was seen that the highest correlation between the two curves was found on average at a lag of -8 trials for the allocentric rules (M=-8.0, SEM=3.78, deviation from zero Wilcoxon one sample test: W(5)=0, p=.043) and -6 trials for the cue-based rules (M=-5.52, SEM=2.29, Wilcoxon one sample test: W(30)=96, p=.015), indicating that the decoded directions precede the real directions over the course of learning (see Fig. 4c).

Furthermore, we looked at the distribution of decoded probabilities while the animal was at the choice point, but had not yet entered any of the goal arms. The distribution of decoded probabilities showed an overall increase of decoded bins of the upcoming choice compared to the alternative choice on the maze (interaction maze bin and prediction category, repeated measures ANOVA: F(9,351)=115.14, p<.001; Fig. 4d), with most notable increases in occurrence of decoded timepoints of the start (post-hoc Bonferroni corrected p-value: p<.001) and end (post-hoc Bonferroni corrected p-value: p<.001) of the arm representing the upcoming choice. To look at the relative contribution of SWR-modulated and unmodulated mPFC neurons, we next built separate decoding models for each subpopulation (see Methods). The differences in decoded distributions between the upcoming and alternative choice were also seen when using the SWR-unmodulated mPFC neurons to build the decoding models (interaction maze bin and prediction category, repeated measures ANOVA: F(9,288)=30.1, p<.001; post-hoc Bonferroni corrected p-values, start of the goal arms: p<.001; end of the goal arms: p<.001), but not when using the SWR-modulated neurons for the decoding (interaction maze bin and prediction category, repeated measures ANOVA: F(9,279)=15.53, p=.135; Fig. 4d). This lack of differentiating the upcoming and alternative choices at the center of the maze was true for both the SWR-inhibited and SWR-excited cell populations (SWR-inhibited, interaction maze bin and prediction category, repeated measures ANOVA: F(9,270)=27.36, p=.193; SWR-excited, repeated measures ANOVA: F(9,270)=17.50, p=.154). This indicates a larger involvement of the SWR-unmodulated neurons in encoding the upcoming choice at the center of the maze.

In order to compare these findings with hippocampal spatial mapping, we decoded spatial activity in the hippocampus and obtained representations of upcoming and alternative choices in the same way as in the mPFC (see methods and Supplementary Fig. S4). Due to more limited multi-unit activity recorded from the hippocampus, we were only able to reliably decode hippocampal activity from N=12 sessions from 3 different animals, but the turning direction could still be predicted reliably based on the non-local representations in this region when the animals were in the process of learning a spatial rule (Supplementary Fig. S4b). To look at the overlap of the decoded representations between the hippocampus and the mPFC, we computed the percentage of overlapping classifications for each session and compared this to shuffled data (i.e., the representations in the mPFC per category of local, upcoming and alternative choices, compared to N=500 times the shuffled decoded data from the hippocampus, see Methods). While the overlap between the hippocampus and mPFC in local representations was significantly higher than the shuffled data (overlap local representations: M=10.10% of decoded time bins, SEM=1.30, shuffled population: M=9.08, SEM=1.13, paired samples t-test: t(10)=3.31, p=.008), no such overlap was seen for the representations of upcoming and alternative choices (overlap alternative choice: M=0.11%, SEM=0.04, shuffled population: M=0.11, SEM=0.04, paired samples t-test: t(10)=0.03, p=.973; overlap upcoming choice: M=0.48%, SEM=0.24, shuffled population: M=0.22, SEM=0.09, paired samples t-test: t(10)=1.54, p=.154; Fig. 4e). For the decoded data based on either the SWR-modulated or SWR-unmodulated neurons in the mPFC, no significant overlap in time was seen with the decoded data from the hippocampus. (Overlap local representations, SWR-modulated [shuffled]: M=7.51%, SEM=1.22 [M=6.99%, SEM=1.00], paired samples t-test: t(10)=1.28, p=.228; SWR-unmodulated [shuffled]: M=7.15%, SEM=1.04 [M=6.48%, SEM=0.82], paired samples t-test: t(10)=2.02, p=.071. Overlap alternative choice: SWR-modulated [shuffled]: M=0.92%, SEM=0.60 [M=0.29%, SEM=0.10], paired samples t-test: t(10)=1.18, p=.266; SWR-unmodulated [shuffled]: M=0.86%, SEM=0.61 [M=0.26%, SEM=0.10], paired samples t-test: t(10)=1.09, p=.302. Overlap upcoming choice: SWR-modulated [shuffled]: M=0.66%, SEM=0.19, [M=0.51%, SEM=0.11], paired samples t-test: t(10)=1.03, p=.325; SWR-unmodulated [shuffled]: M=0.80%, SEM=0.25 [M=0.51%, SEM=0.13], paired samples t-test: t(10)=1.55, p=.151. See Fig. 4e). Again, this was the case for both the SWR-excited and SWR-inhibited cell populations (Overlap upcoming choice, SWR-excited [shuffled]: M=0.25%, SEM=0.07 [M=0.18%, SEM=0.04], paired samples t-test: t(10)=1.59, p=.143; SWR-inhibited [shuffled]: M=0.86%, SEM=0.63 [M=0.27%, SEM=0.12], paired samples t-test: t(10)=1.08, p=.305. Overlap alternative choice, SWR-excited [shuffled]: M=0.61%, SEM=0.17 [M=0.46%, SEM=0.09], paired samples t-test: t(10)=1.12, p=.290; SWR-inhibited [shuffled]: M=0.33%, SEM=0.11 [M=0.29%, SEM=0.09], paired samples t-test: t(10)=0.43, p=.674). Thus, the encoding for alternative and upcoming choices on the maze is largely separated in time between the mPFC and hippocampus.

### Predictive representations in the mPFC are not linked to hippocampal SWRs and theta phase

Finally, we were interested to see whether the non-local representations of the upcoming choice in the mPFC are linked to hippocampal SWRs and theta phase. We obtained a histogram of SWR occurrence in temporal proximity to the start of each segment of non-local representations in the mPFC, and compared this to a shuffle distribution of decoded probabilities (see Methods). Only segments of non-local representations were included that occurred within 2 seconds of a hippocampal SWR. The occurrence of decoded segments at SWR onset were not significantly different from shuffled distributions for both upcoming and alternative choices (occurrence of segments of alternative choice at 0-20 ms from SWR onset, real: M=0.06, SEM=0.01, shuffled: M=0.07, SEM=0.01, Wilcoxon signed-rank test: W(33)=190, p=.382; occurrence of segments of upcoming choice at 0-20 ms from SWR onset, real: M=0.06, SEM=0.01, shuffled: M=0.07, SEM=0.01, Wilcoxon signed-rank test: W(33)=174, p=.147; Fig. 5a). It was further seen that the SWR-unmodulated mPFC neurons had higher normalized firing rates than SWR-modulated neurons during representations of upcoming choice (z-scored firing rates of SWR-excited neurons: M=0.24, SEM=0.07, SWR-inhibited neurons: M=0.06, SEM=0.07, SWR-unmodulated neurons: M=0.43, SEM=0.10, Kruskal-Wallis: H(94)=7.55, p=.023; Fig. 5b). During representations of the alternative choice, the difference was not significant (z-scored firing rates of SWR-excited neurons: M=0.18, SEM=0.13, SWR-inhibited neurons: M=0.14, SEM=0.08, SWR-unmodulated neurons: M=0.37, SEM=0.11, Kruskal-Wallis: H(94)=5.0, p=.082; Fig. 5b).

**Figure 5.**
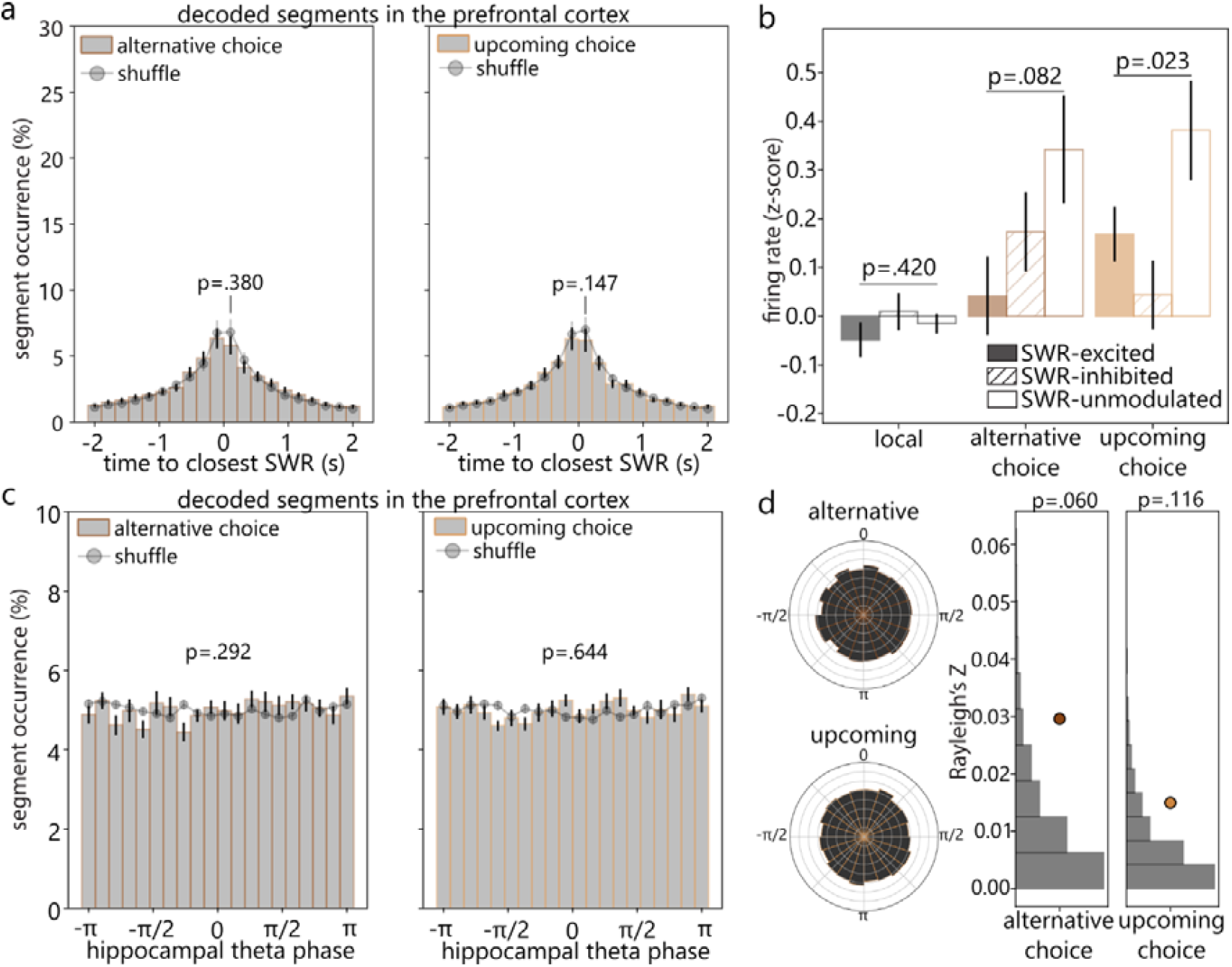
Non-local representations in the mPFC are not linked to hippocampal SWRs and theta phase. **(a)** The occurrence of representations of alternative choice (left) and upcoming choice (right) in the mPFC in temporal proximity to the closest SWR. The p-values indicate the results of a Wilcoxon signed-rank test between real and shuffled data at bin 0-20 ms from SWR onset. **(b)** The firing rates of mPFC neurons that are SWR-excited, SWR-inhibited and SWR-unmodulated, during representations of upcoming and alternative choices, normalized to the average firing rate at the start of the maze. The p-values indicate the results of a Kruskal-Wallis test between the different SWR modulation categories. **(c)** A histogram of the phase of hippocampal theta oscillations at which the start of each representation of upcoming and alternative choices in the mPFC occurred. The p-values indicate the results of a Kolmogorov-Smirnov test for difference in distributions between the representations of alternative and upcoming choices compared to shuffled decoded probabilities. **(d)** Left: circular histograms of the phase locking shown in (c). Right: The phase locking score of representations of upcoming and alternative choices (Rayleigh’s Z, dark and light brown dots) to hippocampal theta oscillations compared to a randomized population (gray histograms). P-values indicate the result of a left-tailed Monte Carlo test of the Rayleigh’s Z compared to the shuffled population.

Similarly, we obtained a histogram of hippocampal theta phase at which each segment of non-local decoding occurred. Only segments of non-local representations that occurred within ‘theta periods’ were included (which were selected based on a running speed of >5cm/s and the absence of SWRs in the hippocampus; see Methods). The distribution of hippocampal theta phase at the onset of representations of upcoming and alternative choices was very homogeneous with no clear bias towards a particular phase of the theta cycle (Kolmogorov-Smirnov difference in distributions, alternative versus shuffled: K=0.25, p=.292; upcoming versus shuffled: K=0.15, p=.644; see Fig. 5c). As such, the average phase-locking scores of upcoming and alternative segments were not statistically significant compared to a randomized population (representations of alternative choice, Rayleigh’s Z=0.03, randomized population M=0.01, CI=[0.00, 0.04], p=.060; representations of upcoming choice, Rayleigh’s Z=0.01, randomized population M=0.01, CI=[0.00, 0.03], p=.116; Fig. 5d). These results thus indicate that the non-local representations in the mPFC are not directly linked in time to hippocampal SWRs and theta phase.

### Predictive representations in the hippocampus are linked to SWRs and theta phase

In order to compare these findings with non-local representations in the hippocampus, we obtained a histogram of SWR occurrence in temporal proximity to the start of each non-local representation in the hippocampus. We found that hippocampal representations of upcoming choice, but not the alternative choice were more time-locked to SWRs than the shuffled probabilities (occurrence of segments of alternative choice at 0-20 ms from SWR onset, real: M=0.12, SEM=0.02, shuffled: M=0.12, SEM=0.01, Wilcoxon signed-rank test: W(14)=59, p=.978; occurrence of segments of upcoming choice at 0-20 ms from SWR onset, real: M=0.18, SEM=0.03, shuffled: M=0.12, SEM=0.01, Wilcoxon signed-rank test: W(14)=12, p=.004; Fig. 6a). Furthermore, interestingly, SWR-excited neurons in the mPFC fired significantly more during hippocampal representations of upcoming choice compared to representations of alternative choice (z-scored firing rates during representations of alternative choice: M=-0.15, SEM=0.05, during representations of upcoming choice: M=0.06, SEM=0.06, Mann-Whitney U test: U(28)=57, p=.023; Fig. 6b). For SWR-inhibited neurons, the opposite effect was seen, where firing rates were higher during hippocampal representations of alternative choice, and decreased during representations of upcoming choice (z-scored firing rates during representations of alternative choice: M=0.15, SEM=0.10, during representations of upcoming choice: M=-0.13, SEM=0.08, Mann-Whitney U test: U(28)=162, p=.042; Fig. 6b). For SWR-unmodulated neurons, no difference was seen in firing rate modulation linked to representations of upcoming and alternative choices in the hippocampus (z-scored firing rates during representations of alternative choice: M=0.04, SEM=0.04, during representations of upcoming choice: M=0.04, SEM=0.03, Mann-Whitney U test: U(28)=100, p=.619; Fig. 6b). This indicates that mPFC neurons that are modulated by hippocampal SWRs, respond selectively to hippocampal representations of upcoming and alternative choices.

**Figure 6.**
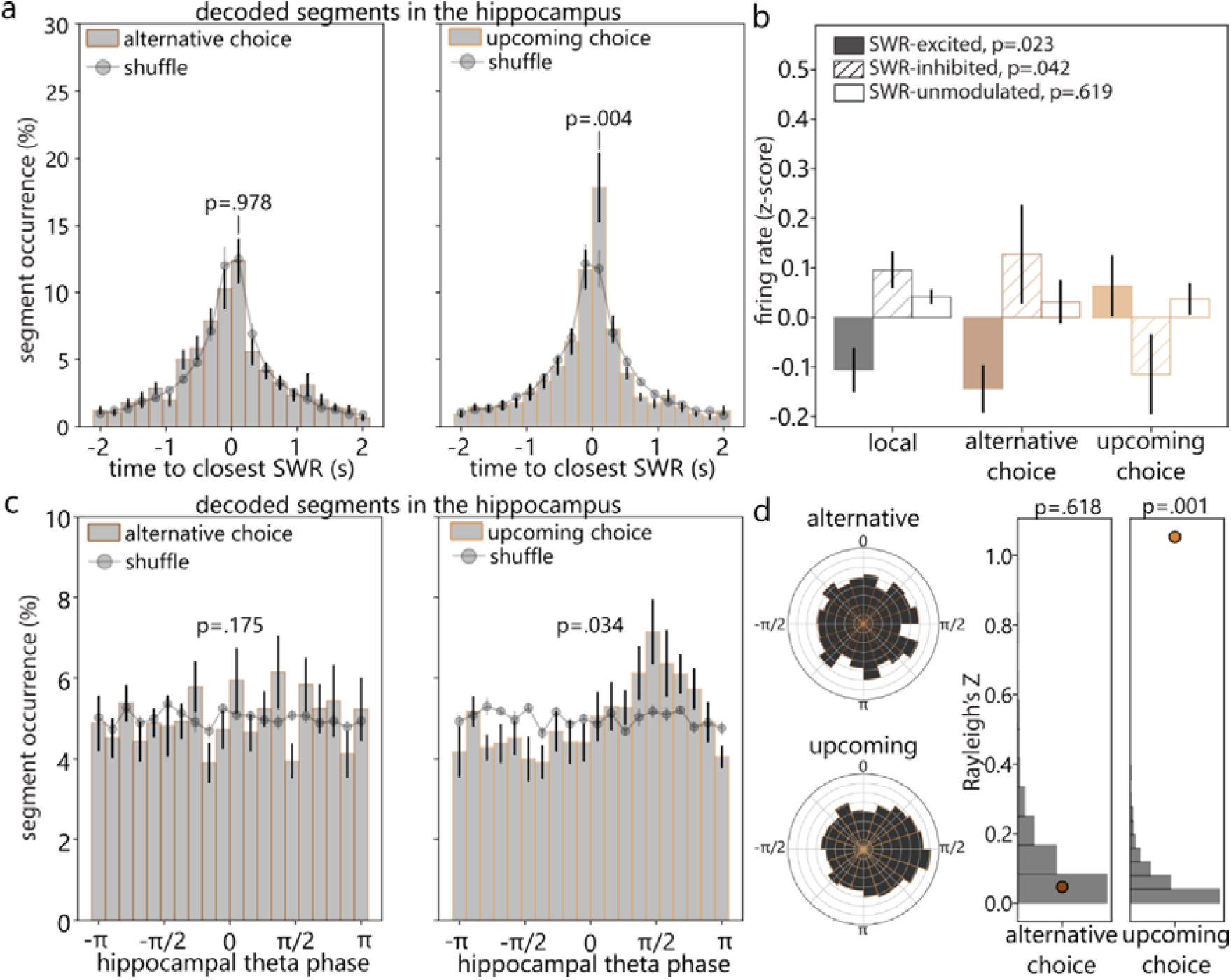
Non-local representations in the hippocampus are linked to SWRs and theta phase. **(a)** The occurrence of representations of alternative choice (left) and representations of upcoming choice (right) in the hippocampus in temporal proximity to the closest SWR. The p-values indicate the results of a Wilcoxon signed-rank test between real and shuffled data at bin 0-20 ms from SWR onset. **(b)** The firing rates of mPFC neurons that are SWR-excited, SWR-inhibited and SWR-unmodulated, during representations of upcoming and alternative choices, normalized to the average firing rate at the start of the maze. The p-values indicate the results of a Mann-Whitney U test for the firing rates during upcoming and alternative representations for each of the SWR modulation categories. **(c)** A histogram of the phase of hippocampal theta oscillations at which the start of each representation of upcoming and alternative choices in the hippocampus occurred. The p-values indicate the results of a Kolmogorov-Smirnov test for difference in distributions between the representations of alternative and upcoming choices compared to shuffled decoded probabilities. **(d)** Left: circular histograms of the phase locking shown in (c). Right: The phase locking score of hippocampal representations of upcoming and alternative choices (Rayleigh’s Z, dark and light brown dots) to hippocampal theta oscillations compared to a randomized population (gray histograms). P-values indicate the result of a left-tailed Monte Carlo test of the Rayleigh’s Z compared to the shuffled population. Due to less multi-unit activity recorded in the hippocampus, fewer sessions were available with good decoding quality, and figure is made with N=12 sessions from 3 animals.

Hippocampal representations of alternative and upcoming choice were differently distributed with respect to the hippocampal theta cycle (Kolmogorov-Smirnov difference in distributions, alternative versus shuffled: K=0.35, p=.175; upcoming versus shuffled: K=0.45, p=.034; Fig. 6c). The representations of the upcoming choice occurred more during the rising phase of the theta cycle (Rayleigh’s Z=1.05, randomized population M=0.06, CI=[0.00, 0.23], p=.001), whereas the representations of alternative choice were more scattered in relation to the hippocampal theta cycle (Rayleigh’s Z=0.05, randomized population M=0.09, CI=[0.00, 0.33], p=.599; Fig. 6d). Together, these results indicate that while mPFC non-local representations are not time-locked to hippocampal SWRs and theta phase, the non-local representations in the hippocampus are.

## DISCUSSION

In this study, we propose two populations of mPFC neurons that are important for spatial learning and decision making. The first is more strongly influenced by hippocampal events, showing firing patterns linked to hippocampal SWRs, hippocampal theta oscillations and hippocampal non-local representations of upcoming choice. A second population of mPFC neurons is less modulated by hippocampal events. These neurons have higher spatial tuning properties, fire more selectively during certain head movements, and fire more during non-local representations in the mPFC that are predictive of the upcoming choice.

We first showed that neurons in the mPFC are modulated by SWRs in the hippocampus, and this modulation was stronger for the ventral mPFC than the dorsal section of the mPFC. The hippocampus and ventral mPFC are connected through several pathways^42^. The connection with the dorsal mPFC, on the other hand, is much less established. One study has reported a monosynaptic prefrontal cortex projection from the ACC to the hippocampus^43^, but no direct projection from the hippocampus to the ACC. The denser connections with the ventral mPFC could explain the stronger SWR modulation seen in the ventral compared to the dorsal mPFC.

Congruent to previous research^12^, we next showed that mPFC neurons that were modulated by hippocampal SWRs, were also phase locked to the hippocampal theta rhythm, suggesting a network of hippocampal spatial mapping and mPFC firing patterns that is co-modulated during both running and resting periods. We showed that these findings hold up for both the dorsal and ventral mPFC, although the preferred phase of the hippocampal theta cycle was different. In the dorsal mPFC, neurons fired more at the peak of the oscillation, while neurons in ventral mPFC fired more just before the trough of the oscillation. An influential paper published in 2009 by Lubenov and Siapas^44^ has shown that theta oscillations are organized topographically across the hippocampus, indicating that a segment of space, rather than a point, is represented across the hippocampus at any given point in time. This concept of a ‘traveling wave’ of theta oscillations might translate to the downstream cortical regions as well, as our data suggests that cells in the mPFC are phase locked to later phases of the hippocampal theta cycle in the more ventral regions.

We further showed that spatial locations could be reliably decoded based on spiking activity in the mPFC, and that the non-local representations at the start of the maze predicted the future choice of the animal. Moreover, following a rule switch, the mPFC decoded probabilities changed a few trials before the behavioral switch, which points to a leading role of the mPFC in spatial rule switching. There was very little overlap between the non-local representations in the mPFC and the hippocampus for the representations of the upcoming choice and alternative choices. This indicates a separation in time between the two regions when engaging in the planning of future trajectories and representing alternative choices.

Previous research has shown that spatial mapping in the hippocampus is linked to theta oscillations during running^45,46^, and spatial reactivations are frequently known to occur during SWRs while resting^6,7,13,31,47,48^. Here we showed several pieces of evidence that suggest that spatial mapping in the mPFC occurs at least partially independently from SWRs and theta phase in the hippocampus. First, neurons in the mPFC that were not modulated by hippocampal SWRs, showed higher spatial tuning scores, head direction selectivity and theta cycle skipping properties. The spatial tuning scores and head direction selectivity indicate that these SWR-unmodulated neurons are important for spatially mapping the environment, and theta cycle skipping properties in the mPFC have in the past been linked to signaling upcoming choice^39,41^. The increased involvement of SWR-unmodulated neurons in these functions was robust and consistent across subregions. Second, non-local representations in the mPFC that were predictive of the animals’ upcoming choice, were not linked to hippocampal SWRs and theta phase. In contrast, non-local representations in the hippocampus showed a much higher phase locking score to hippocampal theta oscillations, and these predictive segments were time-locked to SWR events. Finally, SWR-unmodulated neurons in the mPFC preferentially encoded the upcoming choice over the alternative choice at the center of the maze, and these neurons fired more during segments of upcoming choice in the mPFC than the SWR-modulated neurons. This suggests a role for signaling upcoming choice for those neurons that are not modulated by hippocampal SWRs. In contrast, SWR-modulated neurons were linked to non-local representations in the hippocampus, with firing rate modulations that were homologous to their modulation during SWRs (SWR-excited neurons fired more during hippocampal representations of upcoming choice and SWR-inhibited neurons fired less during hippocampal representations of upcoming choice).

These findings are not directly intuitive, given the running hypothesis that that the hippocampus transfers information to the cortex during SWR events^11^, which could lead one to hypothesize that neurons in the cortex that respond to hippocampal SWRs are responsible for integrating spatial information and creating a spatial map necessary for effective spatial decision making. This hypothesis is supported by several previous findings. For one, a recent paper has shown that spatial maps in several cortical areas become less well defined following hippocampal lesions, indicating that cortical regions rely on input from the hippocampus in order to create a spatial map^49^. A recent paper from our lab further showed that disrupting the mPFC during the 200 ms following SWR onset, slowed learning of a new rule in a spatial alternation task^14^, indicating that the input from the hippocampus during SWRs is indeed necessary for adaptive learning in response to changing reward contingencies. However, none of these findings stipulate that the neurons that respond to hippocampal SWRs should be the same neurons that integrate the spatial map and are involved in establishing a behavioral strategy. Neurons in the mPFC that are modulated by hippocampal SWRs have been shown to be selectively modulated by specific reactivation patterns or replay sequences^13^. Therefore, we hypothesize that the population of neurons that is modulated by SWRs is important for signaling which hippocampal reactivation patterns are important, after which another population of mPFC neurons may alter its spatial map accordingly and signal the adapted behavioral strategy following the external rewards and task demands.

### Suggestions for future research

Together, these findings raise some interesting questions about the function of SWR-modulated and SWR-unmodulated neurons in spatial learning. We hypothesize that SWR-modulated and SWR-unmodulated neurons work together by respectively integrating information from the hippocampus and altering the spatial map necessary for effective spatial decision making. To test this hypothesis, a future study could detect SWRs in the hippocampus and inhibit mPFC activity, while also performing high-density probe recordings in the mPFC to obtain information about single cell firing patterns. If the SWR-linked inhibition of the mPFC alters mPFC spatial coding, by lowering the prediction index of the non-local representations for example, this would support the aforementioned hypothesis. Finally, subregion specific disruptions of the dorsal and ventral mPFC, using optogenetics for example, could test whether there is a differential involvement of the two regions in the spatial decision making process. A task with more structured time points of decision making would be recommended to disentangle the times at which this information is encoded in the different subregions.

## Conclusion

In conclusion, neurons in the mPFC are able to predict future choice on a maze on a trial-to-trial basis and their prediction precedes the behavioral change over the course of learning. This suggests an important role for the mPFC in spatial learning and decision making. Our results further show that SWR-unmodulated neurons are active at key time points in spatial decision making, and timepoints of non-local decoding in the mPFC and hippocampus do not tend to overlap. This indicates that SWR-modulation is likely not the driving factor behind the non-local decoding in the PFC.

## METHODS

### Animals

Six male Long Evans rats were used in this study, weighing between 300 and 400 grams. The animals were allowed at least one week to acclimatize upon arrival, during which time they were housed in pairs and had unlimited access to food and water. During the course of the experiment, the rats were single housed and kept at a mild food restriction schedule, maintaining over 85% of their initial body weight.

### Apparatus

The experiments were conducted in a room of 2.9 by 4.2 meters, with in the center a plus shaped maze that was elevated 50 cm above the ground. The four arms of the maze were 60 cm each, with at the end a platform of 20 by 20 cm holding a reward container, and in the center a piece of 38 by 38 cm.

### Pre-training

Pre-training was performed with a protocol adapted from previous literature^50^, and consisted of four stages during which the animals slowly got acclimatized to the maze and learned to forage for food reward pellets (chocolate covered rice cereal). First, the rats were familiarized with the maze during 5 minutes of free exploration. Next, the day after, 10 reward pellets were spread across the maze and the animals were allowed to forage for 15 minutes. On the third day, three reward pellets were placed in each of the four reward containers and the animal was again allowed to forage for 15 minutes. After this, on all consecutive days, only one reward pellet was placed in each reward container, and the animal had to empty all four reward containers in order for them to be refilled. The maze familiarization was completed when the animal consumed all four rewards at least four times within 15 minutes. The animals took on average 8 days to complete the pre-training.

### Allocentric and cue-based rule learning on the plus maze

Animals had to learn allocentric and cue-based rules in the maze, which were to be deducted through trial and error over the course of multiple days. Each day started with a rest period of 20 minutes. After this, the animal was placed by the experimenter on either the East or the West arm of the maze pseudo-randomly and it had to learn to turn to the North, South, illuminated or non-illuminated arm. The order of the rules was counterbalanced between animals, and the illumination of the arm was always present (also during allocentric learning) and was pseudo-randomly determined for each trial. After arriving at the end of the goal arm, the animal was picked up by the experimenter and placed back in the dark resting box for 30-60 seconds before continuing with the next trial. The training lasted for a maximum of 1.5 hours or 60 trials per day. At the end of the session, animals were placed back in the dark resting box for up to 60 minutes. After reaching the learning criterium (more than 80% of trials correct on three consecutive days), the animals underwent probe implantation surgery (see below). After surgery, the same rule was reinstated until learning criterium. After reinstatement, the rule was switched, either within the same strategy (i.e., rule reversal – from North to South), or to a different strategy (i.e. attentional Set-shift – from North to Illuminated arm), without any indication for the animal. Throughout learning, one of two sounds was played every time the animal passed a sensor halfway through the end arm (at 90 cm along the track). The sounds, a high and low buzz, were associated with correct and incorrect trials. They therefore provided feedback to the animals about whether a reward would be obtained, a few seconds before actually receiving it. Which sound was played to indicate the outcome was counterbalanced across animals but remained constant for each animal during learning. The recordings were completed once the animals had learned all four rules.

### Implant preparation

To record electrophysiological signals from the hippocampus and mPFC simultaneously while the animals were freely moving in the maze, a dual probe implant was used, designed to hold two 1.0 Neuropixels probes (Imec, Leuven, Belgium) at a distance of 6.28 cm^32^. For one of the six animals, there was no probe implanted in the hippocampus, so this animal was excluded from the SWR-related analyses. Prior to implantation, the implant was assembled containing the two probes. The probes were aligned in the stereotaxic frame to be as parallel to each other as possible and perpendicular to the base of the implant. Two holes at the base of the implant were covered in a mixture of bone wax and mineral oil, and the whole surface was cleaned with Baytril (50 mg/ml). The probes were dipped in isopropanol for sterilization, and then in DiI (DIIC18(3) orange/red carbocyanine dye 549/565 nm, VWR, Radnor, Pa, USA) for visualization of the probe tracks later on.

### Surgical procedures

Surgery was performed under full anesthesia, using isoflurane (1-2%) administered via a nose cap and adjusted according to the vital signs of the animal (measured with PhysioSuite monitoring system, Kent Scientific, Torrington, CT, USA). A stereotaxic frame with ear bars kept the head in position and the eyes were covered with eye ointment and aluminum foil for protection. After shaving and disinfecting the area, a midline incision was made from the eyes to the ears, exposing both the lambda and bregma on the skull. Craniotomies were drilled above the right hemisphere at the coordinates 2.0 ML and -3.4 AP for the hippocampus and 0.4 ML and +2.7 AP for the mPFC. After carefully removing the dura in both craniotomies, the probes were inserted with a micromanipulator at a speed of 20 µm/s to a depth of maximum 7 mm.

The implant was anchored with 8 self-tapping bone screws (1.19 × 4.8 mm self-tapping bone screws, 19010-00, Fine Science Tools, Foster City, CA, USA) placed in a semi-circle in the scull around the implant. One of the posterior screws was connected to grounding wire which in turn connected to the ground of both probes. The implant was fixed using C&B Metabond (Parkell, Edgewood, NM, USA) covering all the bone screws and the exposed part of the scull. After this, the isoflurane level was lowered to 0% and the animal was carefully monitored while regaining consciousness. Post-operative care included subcutaneous injections of 0.35 ml Metacam as an analgesic and topical application of antibacterial cream (Picri-Baume) and an anesthetic cream (lidocaine) immediately after surgery and on the three days following.

### Histology

After completing all the training, animals were euthanized using pentobarbital (5ml intraperitoneal injection of Dolethal, 200 mg/ml). Consecutively, they were perfused transcardially with phosphate buffered saline (PBS), followed by a 4% formaldehyde solution. The brain was removed and stored in a cold room (4^°^ C). After 24 hours, it was transferred to a 30% sucrose solution where it was left for at least three days, until it sank to the bottom of the tube. Brains were sectioned using a vibratome (VT1000 S, Leica, Wetzlar, Germany) at 100 µm thickness, after which they were left to dry overnight. Brain sections were stained with a fluorescent Nissl stain (NeuroTrace 530/615 Red Fluorescent Nissl Stain, Thermo Fisher, Waltham, MA, USA), and the probe tracks were visualized with a fluorescent microscope (MVX10, Olympus, Tokyo, Japan). In order to define the subregions of the mPFC, the probe tracks were reconstructed and mapped using a rat brain atlas^51^. To get the most accurate estimation, information was used from the histological images, together with the known insertion depth during surgery, and the electrophysical signatures in the hippocampus (depth at which SWRs occurred). For one animal only the dorsal mPFC was included, and for the other five animals we recorded from both subregions (see Supplementary Fig. S1).

### Data acquisition

Neural data was acquired at 30 kHz using a PXIe acquisition module (National Instruments, Austin, TX, USA) and SpikeGLX software (https://billkarsh.github.io/SpikeGLX/) and video data was acquired at 50 Hz via an overhead camera and recorded with Cheetah software (Neuralynx, Bozeman, MT, USA). An Arduino Uno (Arduino, Turin, Italy) was used to send pulses at fixed intervals but varying durations to the ongoing neural data and video acquisitions, to facilitate the time alignment of the two streams later on (see below).

### Data analysis

#### Position data

Position data was determined using DeepLabCut^52^, a deep neural network algorithm designed for animal behavior tracking. 1084 frames were labelled manually indicating a red and blue LED at the top of the implant, and the tail base of the animal. The model was trained for 500,000 iterations, and the obtained position data was then smoothed, removing any spurious position detections outside of the maze area and position jumps of more than 20 pixels. The positions were linearized in a range from 0 to 240 cm, where positions 0-60 and 120-180 were the two start arms (West and East), and positions 60-120 and 180-240 corresponded to the two goal arms (South and North respectively). For decoding, the position data was binned into 5 cm bins and smoothed with a Gaussian kernel (SD = 5 cm).

#### Time synchronization

A time alignment function was created for each session in order to associate the neural signals with the positions in the maze at which they occurred. The time alignment was based on the duration of pulses sent to both systems (neural- and position data) generated by an Arduino Uno. For each timepoint in the analysis, the corresponding timestamp of the other system was extrapolated using a cross-correlation of the pulse durations.

#### Spike sorting

Data was pre-processed using CatGT (https://billkarsh.github.io/SpikeGLX/#catgt) with a local common average reference (CAR), using the average between rows 2 and 8 from each electrode as a reference. The pre-processed data was then sorted using Kilosort 2 (https://github.com/MouseLand/Kilosort/releases/tag/v2.0), with a detection threshold of 5 and a lower frequency limit of 150 Hz. The output for each session was manually curated using Phy2 (https://github.com/cortex-lab/phy), and clusters (putative single cells) were selected based on their waveform, principal component features and conformity to the refractory period. After manual curation, the selected clusters (N=11657) underwent an objective quality assessment using spike interface^53^, where only the clusters that were compliant with the neural refractory period (an inter-spike-interval violation score of less than 0.1, corresponding to less than 10% of contaminating spikes) were included for further analysis. This step eliminated 2798 clusters. Then, putative fast-spiking interneurons were detected based on the shape of the waveform and the overall firing rate of the neuron (spike half-width < 0.15 ms and firing rate > 17 Hz)^12^, which eliminated an additional 189 clusters. Finally, clusters that showed very low spiking in the SWR modulation windows (less than 50 spikes; N=2727) were also excluded from analysis, as this could lead to spurious SWR modulation classifications. These procedures led to a total of 5943 remaining clusters to include for analyses.

#### SWR detection

For each animal with a probe in the hippocampus (N=5), one channel was selected from the top layer of dorsal CA1. This signal was filtered between 160 and 225 Hertz, and a contrast frequency signal was calculated by subtracting the average power of the neighboring frequency bands (100-140 Hertz and 250-400 Hertz), in order to eliminate potential spurious detections due to bursts of high frequency oscillations that are not restricted to the ripple frequency range (e.g. chewing artefacts). A threshold for detection was determined by taking the median amplitude of the computed contrast frequency signal, plus 9 times the median absolute deviation (MAD). A low threshold of the median amplitude plus 0.5 times the MAD was used for determining the start and end times. SWRs were only included if they were longer than or equal to 30 milliseconds in duration.

#### SWR modulation

Spike rates from sorted neurons in the mPFC were obtained and normalized to the pre-ripple window (from 1 until 0 seconds before SWR onset). For each cluster, SWR modulation was calculated by averaging the spike rates over all the SWRs of that session. Neurons were considered SWR-modulated if the absolute peak of the z-score in the first 200 milliseconds after SWR onset was significantly higher than the shuffled population. The shuffled population was obtained by taking the spike rates of the cluster around all SWRs of that session, shuffling along the time axis, and calculating the peak z-score for each of the N=250 shuffles. Statistical significance was determined by using a Monte-Carlo p-value considering an alpha of .05.

#### Theta phase locking

Theta epochs were selected based on a running speed of more than 5 cm/s and the absence of SWRs in the hippocampus. The theta phase was then calculated by filtering the hippocampal LFP between 9 and 12 Hertz and taking the angle of the Hilbert transformed signal. A Rayleigh’s Z score with associated p-value was calculated for each cluster and theta phase-locking was determined based on the obtained p-value considering an alpha of .05.

#### Spatial tuning

To calculate a spatial tuning score for each sorted cluster, the maze was segmented into bins of 5 cm. To avoid SWR-modulated spiking from biasing the data, spike rates during SWR windows (0 to 200 ms after the onset of SWRs) were not included in this analysis. The spike rate in each maze bin was calculated by taking the number of spikes recorded in that bin divided by the time spent there over the course of a session. Then the spatial tuning score was calculated using a method described in previous literature^54^, where the normalized spike count per maze bin is multiplied by the base-2 logarithm of the normalized spike rate, and the output is summed over all maze bins. This gives a spatial index score expressed in bits per spike, which indicates how much spatial information each spike carries. The spatial information score was only calculated for a cluster if there were at least 50 spikes in the maze during the session (more than 2 times the number of bins).

#### Head direction selectivity

For each position at the start of the maze, the head direction of the animals was determined based on the two LEDs on top of the animals’ head. Head direction angles were divided in 4 equal quadrants based on the orientation of the maze. The head directions towards the goal arms were then categorized in each of three categories: towards future choice or alternative choice, towards North or South and towards left or right. The spike rates during each head direction were calculated for each neuron, and a neuron was labeled directionally selective if the firing rate was significantly higher during one of the head directions compared to its opposite counterpart (e.g. significantly higher during head directions towards the upcoming choice compared to the alternative choice). Statistical significance was determined by comparing the spike rates to a population of 250 shuffles, where the spike rates per trial were swapped randomly. A directional selectivity index was then calculated by subtracting the overall firing rate during one direction from the firing rate during the other direction and dividing by the sum of both firing rates for each direction category. Thus, this resulted in three scores between -1 (the neuron fired only during head directions towards the alternative choice, towards the South arm, or towards right respectively) and 1 (the neuron fired only during head direction towards the upcoming choice, towards the North arm, or towards left). Head directions in the other two quadrants (forward and backwards) were not taken into account. Since the head direction selectivity was stronger in the SWR-unmodulated neurons for each of the six directions (see Supplementary Fig. S3), the overall absolute directional selectivity index was calculated by obtaining the absolute maximum score out of each of the three categories per neuron.

#### Theta cycle skipping

To determine the theta cycle skipping properties for each cluster, spikes were extracted from periods where the animals were at the start platform, arm and center of the maze. For each of the four possible trajectories on the maze (East-North, East-South, West-North, West-South), an autocorrelation of spike times was obtained to determine the rhythmicity of firing from one spike to the next. The average histogram of spike times was smoothed, and two peaks were extracted: one at around 125 ms and one at 250 ms, corresponding to one and two theta cycles of 8 Hz. The theta cycle skipping index was calculated by subtracting the height of the first peak from the height of the second and normalizing to the maximum height of both peaks. The highest cycle skipping index was selected out of the autocorrelations from the four trajectories. The obtained theta cycle skipping index was then compared to a population of N=250 shuffled scores, which were obtained by a shuffling procedure designed to shuffle the spike times while keeping the existing periodicity of the theta cycle (125 ms) intact.

#### Decoding

To decode spatial information from the hippocampus and mPFC, spikes were detected from the CatGT pre-processed data using an amplitude threshold of -50 µV on the high frequency (300-6000 Hertz) filtered signal. Five features were obtained from each detected spike: depth of the channel, spike amplitude, slope of the waveform, height of the positive peak and latency to the positive peak. A Bayesian decoding algorithm was built, associating the spike features with the position in the maze at which they occurred^55^. Periods where the running speed was less than 15 cm/s were excluded from the model. A median cross-validated decoding error of less than 20 cm was used as an inclusion criterium, which resulted in 40 sessions from 5 different animals for the mPFC decoding and 12 sessions from 3 different animals for the decoding of the data from the hippocampus. For the decoding models based on only SWR-modulated or SWR-unmodulated neurons, the spike times and amplitudes were selected for each neuron that was classified as SWR-unmodulated or SWR-modulated (as well as SWR-excited and SWR-inhibited separately), after which the decoding model was built in the same way as the mPFC decoding, for the same 40 sessions.

#### Prediction index

To predict turning directions based on decoded data, trials were decoded with a bin of 50 ms and 80% overlap, such that every 130 ms was consecutively decoded, corresponding to roughly one theta cycle (at 8 Hz). The decoded probabilities were summed for each maze segment (the two start arms, the center and the two goal arms), and segments of local representations, upcoming choice and alternative choice were obtained by grouping probabilities together in time that had a summed probability of more than 0.5 per maze segment. Segments that were 50 milliseconds apart were joined together. The prediction index per trial was then calculated by subtracting the summed probabilities of the non-taken goal arm from the summed probabilities of the taken goal arm and normalizing to the total summed probabilities of both categories. In this way, a score of -1 corresponds to a trial that only contains probabilities of the non-taken goal arm, and a score of 1 corresponds to a trial where all the non-local probabilities are of the taken goal arm. The prediction indices were separated by whether the spatial rule had or had not been learned, and which spatial rule was in place at the time (allocentric or cue-based). For determining the statistical significance of the prediction index, the posterior probabilities were randomized by shuffling the positions in the two goal arms for each time bin in the start, arm and center of the maze. The prediction index was calculated in the same way for the shuffled data as for the real posterior probabilities, and statistical significance was determined by comparing the real and shuffled data.

#### Prediction accuracy versus learning

To determine whether the prediction accuracy per trial was leading or lagging with respect to the behavior, learning curves were obtained with a moving average of N=10, and overlaid with curves made from the predicted direction based on the non-local representations in the mPFC. The correlation between the two curves (Pearson’s r) was calculated for each trial lag of -N/2 to N/2 (N denotes the total number of trials), such that a peak of correlations at a negative lag indicates that the real direction is lagging behind the predicted direction and a peak of correlations at a positive lag indicates that the real direction is leading the predicted direction.

#### Overlap decoding mPFC versus hippocampus

To assess the overlap of decoding between the hippocampus and the mPFC, each decoded time bin (of 50 ms and 80% overlap), was classified as ‘local’ (the same arm or reward platform as the animal is physically present), ‘alternative choice’ (the arm or reward platform that the animal does not end up visiting on this trial), ‘upcoming choice’ (the arm or reward platform that the animal will visit), or ‘other’ (the other start arm, the center of the maze or low decoding probabilities, i.e. <0.5, for all of the categories). Then the number of overlapping classifications was counted and divided by the total number of decoded time bins of the session and multiplied by 100 to obtain the percentage of overlap. For each session, the classifications from the hippocampus were shuffled 500 times using a rolling method over the position axis. The mean overlap of the shuffled data was compared to the true overlap per session using a paired samples t-test. This procedure was repeated for the decoded data from all mPFC neurons, and for the SWR-unmodulated, SWR-modulated, SWR-excited and SWR-inhibited decoded data separately.

#### Time-locking to hippocampal SWRs and theta phase

To determine the time locking of segments of upcoming and alternative choice in the mPFC and hippocampus to hippocampal SWRs, each decoded segment of alternative and upcoming choice (any number of consecutive time bins where the decoded probabilities of the alternative and upcoming choice respectively were more than 0.5), was assessed for its temporal proximity to hippocampal SWRs. Only non-local segments that occurred within 2 seconds of a SWR were taken into account for this analysis, and only the SWR that occurred closest in time to the non-local segment was included. The histogram of temporal proximity was then normalized to the total number of SWRs per session, and compared to the temporal proximity of shuffled posterior probabilities (randomly rolled over the position axis for each decoded time bin). To determine the phase locking to hippocampal theta, theta epochs were selected as described above (based on a running speed of more than 5 cm/s and the absence of SWRs in the hippocampus), and the theta phase of each decoded bin was determined using a band-pass filter of 9-12 Hertz, and obtaining the angle of the Hilbert transform of this signal. Then, the histogram of theta phases was obtained for the segments of alternative and upcoming choice separately and normalized to the number of segments included. To determine statistical significance, the histograms were compared to shuffled posterior probabilities using a Kolmogorov-Smirnov test.

#### Statistics

All statistical tests were performed using the SciPy library for python^56^. All comparisons were done on the means per session, so that sessions with a very high number of neurons in a certain subregion did not skew the results for that subregion. For every comparison, the assumptions of normality and heteroscedasticity were tested and non-parametric tests were chosen if the assumptions were violated. The difference between firing rates of SWR-modulated and SWR-unmodulated neurons was calculated using an ANOVA or Kruskall-Wallis test (non-parametric). For comparisons between two groups, independent samples t-tests or Mann-Whitney U tests (non-parametric) were used.

## Acknowledgements

We thank Frederic Michon, Cagatay Aydin and Rik van Daal for their contributions in the set-up for chronic Neuropixels probes recordings in freely moving animals and Marine Guyot for her software contributions. F.K is funded by Research Foundation Flanders (FWO), Belgium under grant number G0A5422N, and grant number G077321N and KU Leuven, Belgium C1 grant C14/17/042.

## Author contributions

Conceptualization, H.d.B. and F.K.; Methodology, H.d.B. and F.K.; Software, F.K.; Formal analysis, H.d.B, and F.K.; Investigation, H.d.B; Writing - Original draft, H.d.B, and F.K.; Writing - review and editing, H.d.B., and F.K.; Visualization, H.d.B. and F.K.; Supervision, F.K.; Funding Acquisition, F.K.

## Competing interests

The authors declare no competing interests

## Materials & Correspondence

Correspondence and request for materials can be addressed to Hanna den Bakker.

## Data Availability

The data that support the findings of this study are available from the corresponding author upon reasonable request

## Notes

### Competing Interest Statement

The authors have declared no competing interest.

### Summary of Updates

We have updated our manuscript on three main points of feedback from our reviewers. (1) We have performed additional analyses to further disentangle the contributions of SWR-modulated and SWR-unmodulated PFC neurons to the non-local representations of upcoming choice. (2) We have added a statistical analysis to look at the time-locking of non-local representations to hippocampal SWRs and theta phase. (3) We have rewritten the text to better emphasize our novel contributions and clearly distinguish new findings from confirmatory observations.

