## supplementary figures for "Neurons in the medial prefrontal cortex are involved in spatial tuning and signaling upcoming choice independently from hippocampal sharp-wave ripples"

| Animal ID | N sessions | N clusters PFC | | N clusters HPC |
| --- | --- | --- | --- | --- |
|  |  | dorsal | ventral |  |
| A | 8 | [81,117] | [23,46] | *no HPC probe* |
| C | 13 | [15,41] | [15,74] | [0,6] |
| D | 14 | [0,61] | *not reached* | [0,18] |
| E | 14 | [1,94] | [124,203] | [0,6] |
| F | 9 | [43,111] | [0,14] | [0,14] |
| I | 13 | [113,184] | [13,35] | [26,77] |

***Supplementary Table 1.*** *Overview of the number of sessions per animal and the range [minimum, maximum] of clusters per session per (sub)region. One animal (A) had no probe in the hippocampus, and one animal (D) was implanted too posterior, so only the dorsal mPFC was reached.*

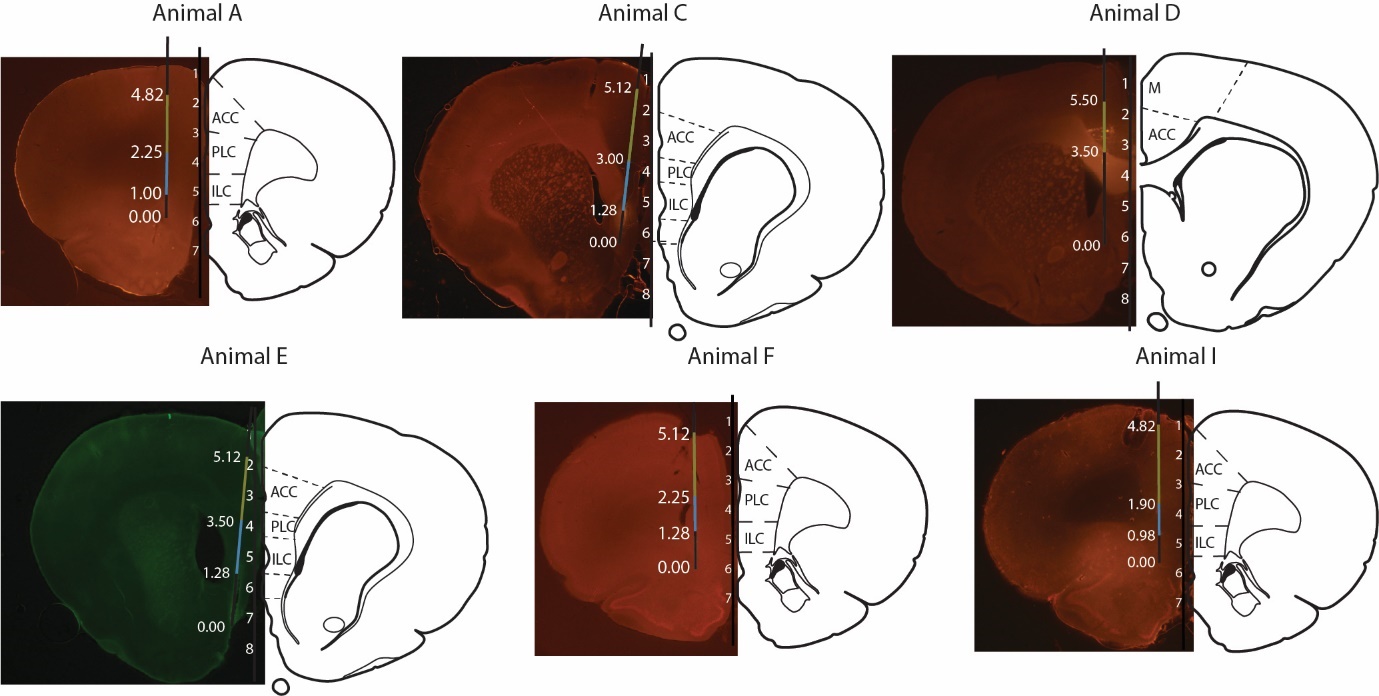

***Supplementary Figure S1. Probe mapping in the mPFC.*** *Probe tracks were reconstructed for all six animals based on the histology, implantation depth and electrophysiological signatures. The numbers indicate the distance from the tip which we used to differentiate the dorsal (in olive) the ventral (in blue) mPFC.*

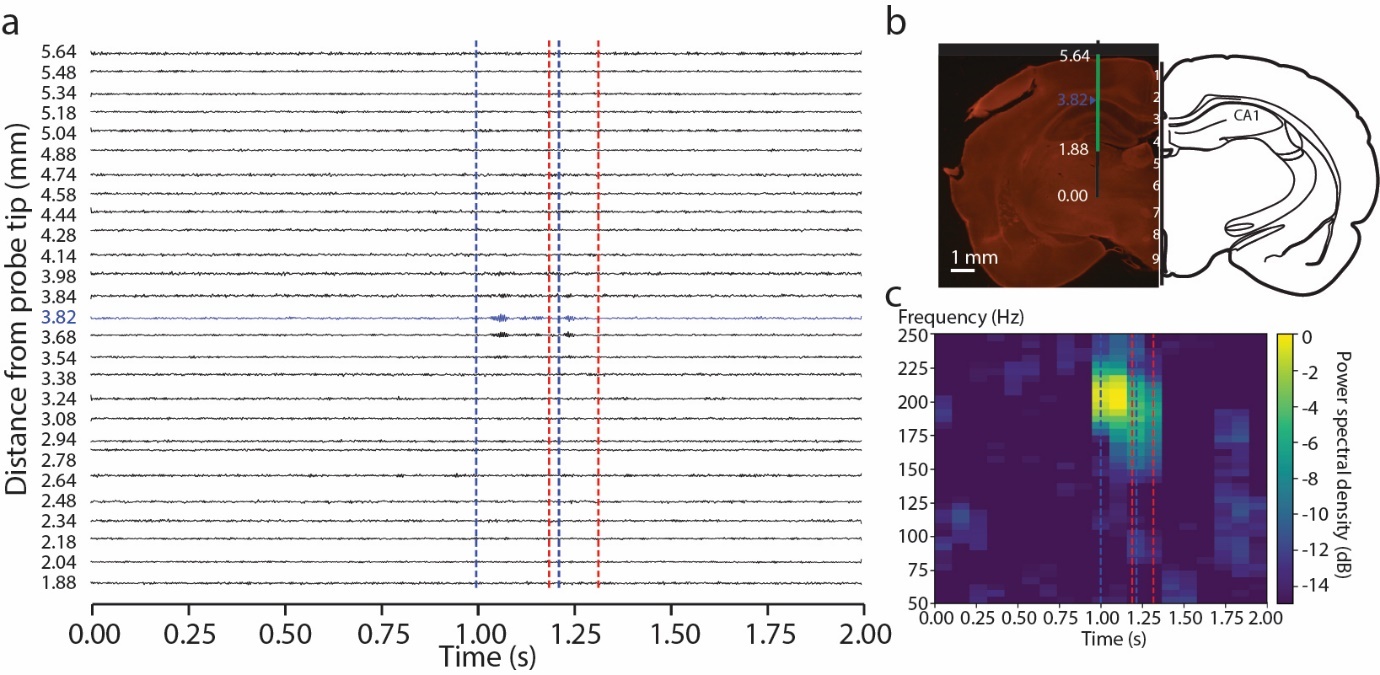

***Supplementary Figure S2. SWR detection.*** *(****a****)* *Two seconds of data from one example animal with the channel used for SWR detection indicated in blue, and the start (blue) and end (red) of two detected ripples indicated with dashed lines. (****b****) A reconstructed probe track of the hippocampus for the same animal as used for panel (a) with the numbers indicating the distance from the probe tip (in mm) and the channel used for SWR detection indicated in blue****.*** *(****c****) A spectrogram of the two seconds of data from the blue channel shown in panel (a), for the frequencies between 50 and 120 Hertz.*

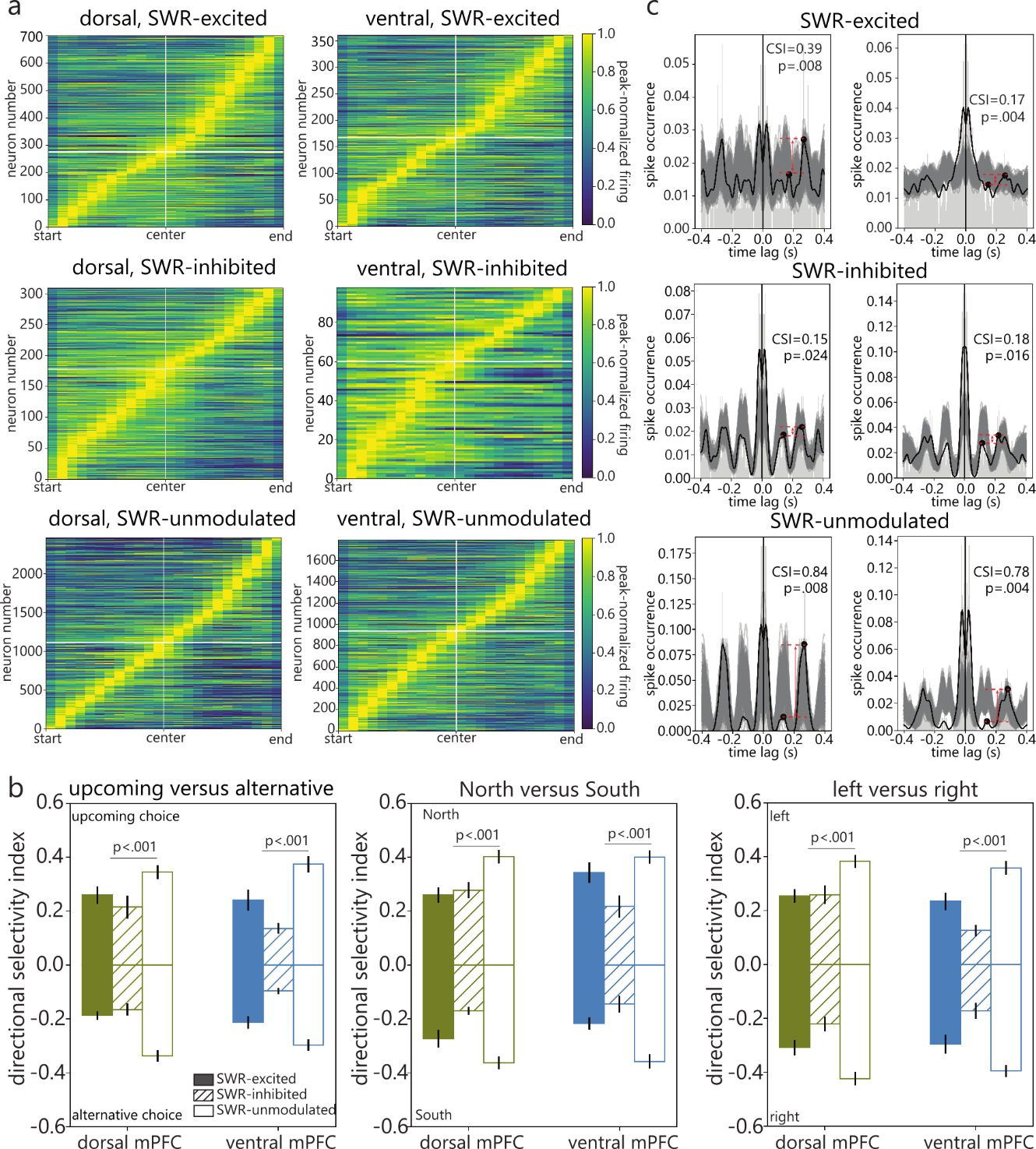

***Supplementary Figure S3. Characterization of spatial tuning, theta cycle skipping and head direction selectivity in SWR-modulated and unmodulated neurons.*** *(****a****) The peak normalized firing rates on the maze sorted for location of the peak rate. The white horizontal lines indicate the border between clusters with a peak rate at the start of the maze, and clusters with a peak rate at the end of the maze. (****b****) Examples of theta cycle skipping cells that were SWR-excited (top row), SWR-inhibited (middle row) and SWR-unmodulated (bottom row). The randomized data is shown in grey, and in black is the smoothed signal, with the peaks at 125 ms and 250 ms (one and two theta cycles) indicated with markers. (****c****) The directional selectivity index (spike rates of neurons during one head direction normalized to the spike rates of both head directions), for head directions that were towards the upcoming choice or the alternative choice (left panel), towards the North or South arm (middle panel) and towards the left or right arm (right panel), separated for neurons that were SWR-excited (solid fill), SWR-inhibited (dashed fill) and SWR-unmodulated (no fill).*

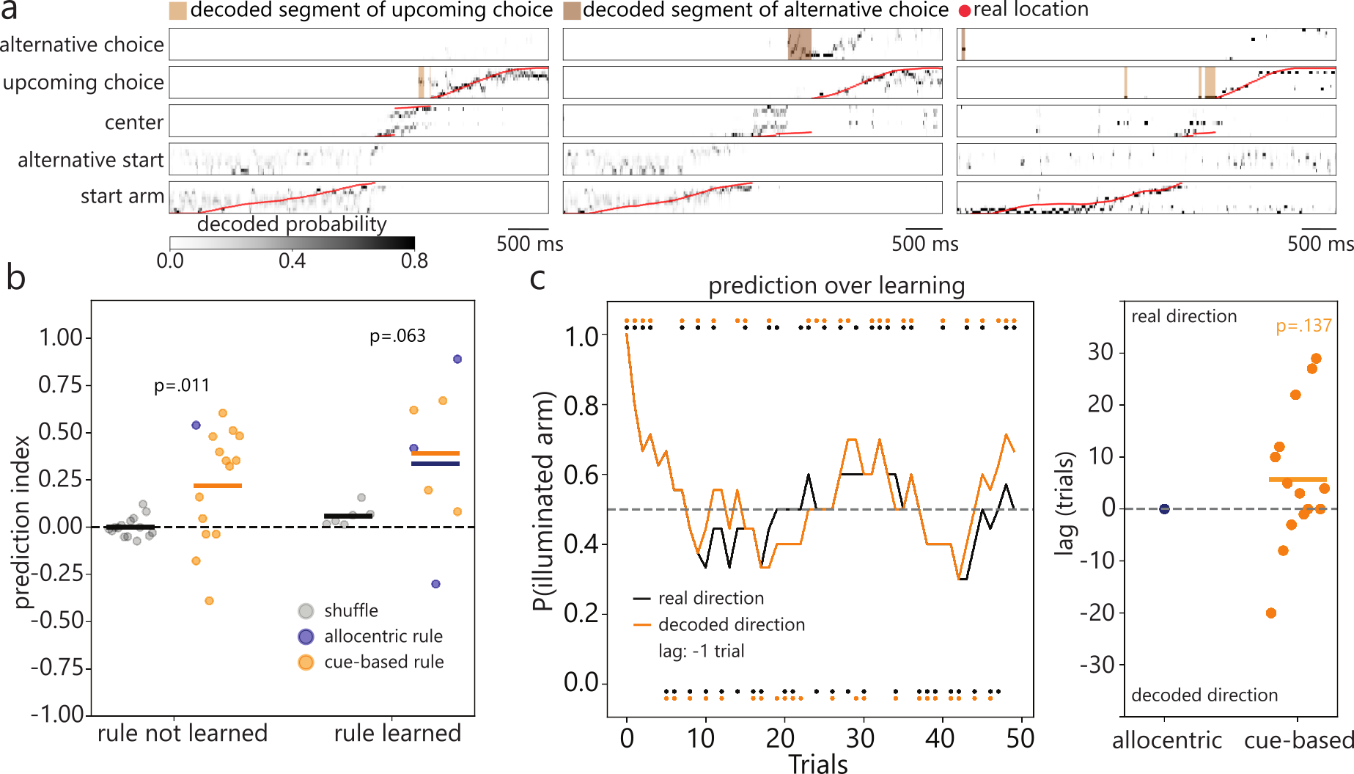

***Supplementary Figure S4. Non-local representations in the hippocampus are predictive of the animals’ upcoming choice.*** *(****a****) Three examples of trials where the decoded positions from the hippocampus (grey scale) and real positions (red) are shown per maze segment. Predictive and non-predictive segments are shown in light and dark brown spans at the top. (****b****) The prediction index per session based on the decoded non-local representations while the animal was at the start of the maze, separated for when the rule was not yet learned and when the rule was learned, and for the allocentric (navy) and cue-based (orange) rules. Shown in light grey is the prediction accuracy based on the shuffled decoded posteriors, and p-values indicate the result of a paired samples t-test between the real and shuffled predictions. (c) Left panel: an example behavioral response curve (in black) where the rule had been changed from the illuminated arm to the dark arm, and the predicted directions based on the non-local representations overlayed (orange). Right panel: the lag (in number of trials) between the real direction and decoded direction per session for allocentric (navy) and cue-based (orange) strategies. The p-value indicates the result of a one sample t-test to test the deviation from zero.*
